# Metagenomics to Metabolomics: Integrating Genomic Insights for Mulberry Crop Protection and Enhancement

**DOI:** 10.1101/2024.05.12.593770

**Authors:** Vidya Niranjan, C Lavanya, H V Pooja, Shreya Satyanarayan Bhat, Spoorthi R Kulkarni

**Affiliations:** Department of Biotechnology, RV College of Engineering, Bangalore, Karnataka, India), Visveswaraya Technological University, Belagavi, 590018; Department of Biotechnology, RV College of Engineering, Bangalore, Karnataka, India., Visveswaraya Technological University, Belagavi, 590018; Department of Biotechnology, RV College of Engineering, Bangalore, Karnataka, India. Visveswaraya Technological University, Belagavi, 590018

**Keywords:** Morus indica, Metagenomics, Metabolites, Biofertilizers, Biopesticides

## Abstract

Indian Mulberry (*Morus indica*) is vital in sericulture, with their leaves serving as the primary food source for silkworms. The soil microbiome surrounding mulberry trees plays a pivotal role in nutrient cycling and ecosystem functioning. This study utilizes metagenomic analysis to explore the taxonomic diversity and functional potential of microbial communities in mulberry soil, particularly focusing on the rhizosphere. Key bacterial species such as *Pseudomonas, Frankia, Azosipirulum* are identified, highlighting their importance in mulberry health and disease dynamics. The abundance distribution of these bacterial populations reveals significant trends, offering insights into mulberry agroecosystem microbial ecology. Understanding garden soil-derived microbial consortia provides a foundation for exploring their role in nutrient cycling and plant health. The study reveals the intricate web of interactions between mulberry and their surrounding soil microbiota and identification of metabolites. Leveraging high-throughput sequencing and bioinformatics, the research identifies potential metabolites as biofertilizers and biopesticides, aiming to improve agricultural sustainability. The findings underscore the critical role of soil microbes in maintaining soil fertility, supporting plant health, and enhancing ecosystem resilience. Despite limitations and gaps, the study contributes to advancing eco-friendly agricultural practices and promoting soil health in mulberry cultivation. Ultimately, the research transcends the laboratory, resonating with stakeholders as it unravels the genetic blueprints of soil life and sows’ seeds of sustainable progress.

## BACKGROUND

Genetic variations in four Mulberry genotypes: Thailand Male (notable for high leaf quality), Assam Bola (root rot resistance), S1 (drought resilience), and Punjab Local (nitrogen utilization). Through SNP and SSR markers, approximately 50 million SNPs were identified, aiding in the development of diagnostic tools and genetic linkage maps. Development of Consensus helped in scaling up the SNPs and finding the best markers. Phenotype-genotype and phenotype-phenotype correlations reveal insights into genetic mechanisms underlying agronomic traits. The research aims to enhance Mulberry agriculture by identifying genetic markers and elucidating genetic linkages, ultimately improving economic viability and environmental sustainability in the sericulture industry. Molecular tools such as SSRs, SNPs, and genetic linkage mapping offer insights into genetic diversity and population structure, facilitating selective breeding efforts. Integration of gene annotation, QTL mapping, and gene expression analysis provides understanding of key traits and pathways, enhancing crop productivity and resilience to environmental stresses. This comprehensive genomic information underscores Mulberry’s significance in agriculture and industry while promoting sustainable development practices. Further, to add value to the study an attempt has been made to extend the studies to Metagenome and metabolomic level to identify the potential biofertilizers and biopesticides for mulberry crop protection.

## INTRODUCTION

Mulberry (Morus spp.) are economically significant plants cultivated globally for their leaves, which serve as the primary food source for silkworms in sericulture. The soil microbiome associated with mulberry trees plays a crucial role in nutrient cycling, plant health, and overall ecosystem functioning [1]. By conducting a metagenomics analysis of mulberry soil, researchers can gain valuable insights into the taxonomic diversity and functional potential of the microbial communities inhabiting this important agroecosystem. The metagenomic sequencing of mulberry samples provides a comprehensive view of the bacterial diversity in both healthy and diseased mulberry plants, particularly focusing on the rhizosphere soil. The identification of key bacterial species, such as *Pseudomonas, Frankia, and Azosipirulum*, highlights the importance of these microbes in mulberry health and disease dynamics.[2] The abundance distribution of these bacterial populations reveals significant trends, with implications for understanding the microbial ecology of mulberry agroecosystems.[3]The metagenomic analysis of garden soil-derived microbial consortia offers valuable insights into the relative abundance of soil microbial communities, This broad survey of soil microbial diversity underscores the complex interactions between microorganisms and their environment, including the soil surrounding mulberry [4]. Understanding the microbial consortia present in garden soil provides a foundation for exploring the role of these microbes in nutrient cycling, plant health, and ecosystem functioning. In essence, mulberry metagenomics research represents a cutting-edge approach to unravelling the intricate web of interactions between mulberry trees and the microbial communities in their surrounding soil. By leveraging high-throughput sequencing technologies and advanced bioinformatics tools, scientists can uncover the hidden microbial diversity, functional potential, and ecological significance of these microbial communities in mulberry agroecosystems.[5] Soil microbes play a fundamental and multifaceted role in the functioning of terrestrial ecosystems, nutrient cycling, and plant health. These microscopic organisms are essential drivers of various biological transformations and the cycling of crucial nutrients, such as carbon (C), nitrogen (N), phosphorus (P), and sulfur (S), which are vital for sustaining soil quality and supporting plant growth.[6] In terms of ecosystem functioning, soil microbes are central to the decomposition of organic matter, releasing plant-available nutrients through their metabolic activities.[7]They also play a pivotal role in carbon cycling, influencing soil structure, and sustainability, and regulating carbon dynamics in the environment through processes like respiration and methanogenesis. Regarding nutrient cycling, soil microbes are involved in key processes like nitrification and denitrification, determining the availability of nitrogen for plants and influencing soil acidification. Similarly, they mediate the transformation and availability of phosphorus and sulfur, ensuring the efficient recycling of these essential elements in the soil ecosystem.[8] Furthermore, soil microbes enhance plant mineral nutrition by improving nutrient availability and uptake through symbiotic associations, such as mycorrhizal fungi and nitrogen-fixing bacteria.[9] The intricate interactions between plants and their associated microbiome in the rhizosphere maximize the nutritional benefits for plant growth and development. Collectively, the multifaceted roles of soil microbes underscore their critical importance in maintaining soil fertility, supporting plant health, and sustaining the overall resilience and productivity of terrestrial ecosystems. The use of biofertilizers containing microbes and its metabolites like their metabolities such as sidophore and geosmine, poluketide, terpene derived from *Pseudomonas, Azospirillum*, and *Frankia* offers a range of potential benefits for sustainable agriculture. *Azospirillum*, a free-living nitrogen-fixing bacteria, can improve shoot and root development, as well as increase the rate of water and mineral uptake by plant roots, contributing to enhanced nutrient availability. *Frankia*, a symbiotic nitrogen-fixing actinobacteria, forms root nodules in non-leguminous plants, helping to fix atmospheric nitrogen and provide it directly to the plants. Additionally, *Pseudomonas* is a phosphate-transforming microorganism that can solubilize inorganic phosphates,[10,11,12] making them more readily available for plant uptake. Beyond their roles in nutrient cycling, these microorganisms can also produce plant growth-promoting substances, such as hormones, vitamins, and amino acids, which can stimulate plant growth and development. Furthermore, the use of these biofertilizers can help plants become more tolerant to various environmental stresses, including drought, salinity, and pathogens. Ultimately, the incorporation of *Pseudomonas, Azospirillum, and Frankia* into biofertilizers can contribute to the overall improvement of soil fertility, structure, and microbial diversity, leading to a more sustainable and productive agroecosystem.[12,14,15]

The current knowledge and application of soil microbiomes and biofertilizers face several limitations and gaps. Market constraints include low product efficacy under unfavorable conditions and the short shelf life of microorganisms, impacting the overall effectiveness of biofertilizers.[16] Environmental factors such as temperature, moisture, pH, and competition with native soil microbiota play crucial roles in determining the efficacy of these products. Concerns also arise regarding the potential introduction of antibiotic-resistance genes (ARGs) into the soil environment through microbial biofertilizers.[17,18] Additionally, the biological restrictions associated with specific microbial strains, including their interactions with existing soil microflora and macrofauna, can limit the performance of biofertilizers.[19] Addressing these limitations is essential for improving the reliability and effectiveness of biofertilizer products in sustainable agriculture, emphasizing the need for further research and development in this field.[20] Our Objectives aim to overcome the limitations and constraints associated with current biofertilizer products and harness the potential of soil microbial diversity for sustainable agriculture. Motivated by the identified challenges such as low product efficacy under unfavorable conditions, short shelf life of microorganisms, potential introduction of antibiotic resistance genes, biological restrictions of specific microbial strains, and environmental factors affecting microbial survival in soil, this research seeks to develop more effective and reliable biofertilizers.[21] By leveraging the insights gained from the metagenomic analysis of mulberry soil, the study will identify key microbial candidates for biofertilizer development. Additionally, the customized database will serve as a valuable resource, allowing researchers to quickly identify if their metagenomic samples contain one or more of the prominent microbes useful for making biofertilizers. This approach will advance eco-friendly agricultural practices, promote soil health, and enhance ecosystem resilience in sustainable mulberry cultivation. The motivation behind researching mulberry soil metagenomics and its microbial inhabitants lies in the profound impact these microorganisms have on agricultural sustainability, soil health, and ecosystem resilience.[22] Understanding the genomic diversity of soil microbes can lead to the identification of potential biofertilizers that enhance soil fertility, promote plant growth, and reduce reliance on chemical fertilizers, revolutionizing sustainable agriculture. Additionally, harnessing microbial communities can aid in soil restoration, erosion control, and carbon sequestration, benefiting both agriculture and the environment [23,24] our findings guide strategies for carbon sequestration and emission reduction, contributing to global climate resilience by managing microbial communities. In summary, our research transcends the laboratory; it resonates with farmers, policymakers, and environmentalists alike, as we unravel the genetic blueprints of soil life and sow seeds of sustainable progress.[25] Our methodology involves a comprehensive approach to analyzing the mulberry soil metagenome, starting with shotgun sequencing to provide a holistic view of all microbial species present. This is followed by focusing on the rhizosphere, the soil surrounding plant roots, to explore specific microbial communities associated with plant health. We also create a comprehensive database of bacterial genomes to facilitate taxonomic classification and functional annotation. Additionally, we quantify microbial abundance at the taxonomic level to provide insights into community structure and guide our focus toward key microbial players. We identify specific microbial taxa with biofertilizer or pesticide potential and review existing research on key microbial species such as *Pseudomonas, Frankia*, and *Azospirillum* to inform our understanding and experimental design. To ensure accurate classification, we curate a high-quality set of reference genomes and create species-specific databases to improve taxonomic assignment accuracy. We link metagenomic reads to specific soil samples to provide spatial context and understand microbial function and interactions. Functional annotation of genes provides insights into microbial capabilities, guiding biofertilizer and pesticide development. Finally, we apply metagenomic insights to create targeted biofertilizers and eco-friendly pesticides, offering practical solutions for sustainable agriculture and soil management.

## 3.0 METHODOLOGY IMPLEMENTED

**Figure 1:**
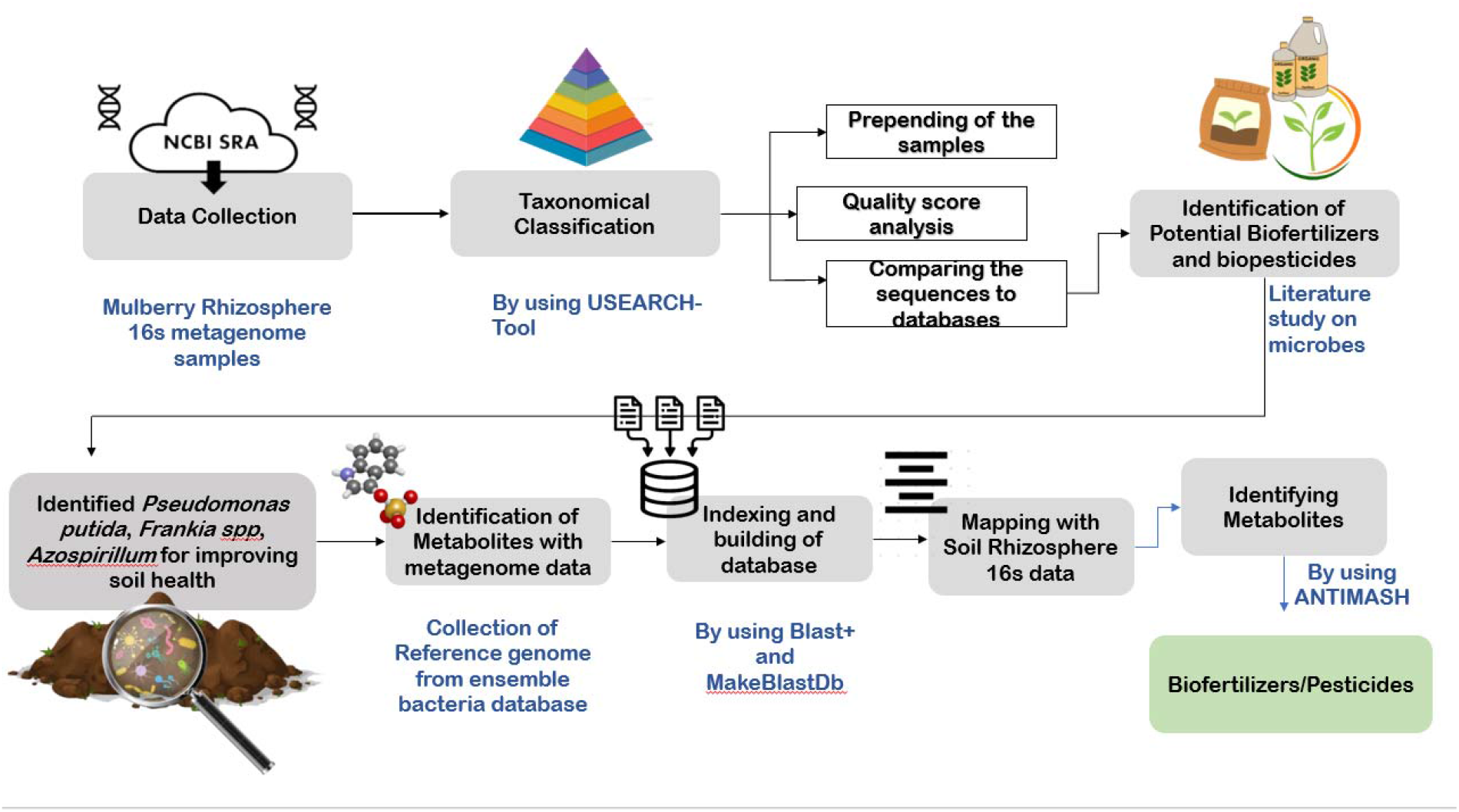
Overview of the Methodology Implemented.

### 3.1 Collection of metagenome samples

The Mulberry Rhizosphere 16s metagenomic samples were collected from NCBI SRA.

The sample shows the following fields:

*Instrument:* Illumina MiSeq

*Strategy:* AMPLICON

*Source:* METAGENOMIC

*Selection:* PCR

*Layout:* PAIRED

### 3.2 TAXONOMICAL CLASSIFICATION OF MICROBES

USEARCH-tool was performed to obtain the taxonomical classification and OTU clustering table. The following is the USEARCH algorithm: USEARCH reads a file containing DNA sequences and sorts them in non-increasing length order. The first step is to combine the fastq files with all of the reads to determine which sample each read belongs to, USEARCH enables us to relabel the reads by prepending the sample name, allowing us to determine which read belongs to which sequence. After that, all combined files are merged to form the merged fasta file. To obtain high-quality read sequences, the reads are trimmed and filtered, yielding the filtered fasta file. De-replication is used to avoid analysing the same sequence twice. De-replication locates a set of distinct sequences in an input file. Sequences are compared letter by letter and must be identical along their entire length (substrings do not match because case is ignored, an upper-case letter will match a lower-case letter). The sintax command predicts taxonomy for query sequences in FASTA or FASTQ format using the SINTAX algorithm. Taxonomy assignment is accomplished by comparing our sequences to the many databases that are available.

### 3.3 IDENTIFICATION OF POTENTIAL BIOFERTLIZERS AND BIOPESTICIDES

The analysis was performed to identify the microbes and their beneficial role in enhancing the crop and additionally, a thorough literature study was performed in order to find the potential microbes which can behave as biofertilizer and a biopesticide. From the literature it was evident that *Pseudomonas putida, Frankia spp, Azospirillum* are important and play a significant role in improving soil health.

### 3.4 IDENTIFICATION OF METABOLITES WITH THE METAGENOME DATA

#### 3.4.1 Collection of Reference Genomes

The reference genomes of *Frankia casuarinae, Azospirullum brasilense, Frankia torreyi* and *Frankia alni* were collected from ensemble bacteria database in fasta format.

#### 3.4.2 INDEXING AND BUILDING OF DATABASE

Latest version of BLAST + was installed and the retrieved sequences was indexed and species wise database was created with the MakeBlastDB option of BLAST+

#### 3.4.3 FINDING THE REGIONS OF SIMILARITY

The 16s metagenome samples were subjected to BLASTn analysis with the database created in the previous step, with this specific gene segment of the sample getting mapped to the reference genome with the identity percent was revealed with the analysis.

#### 3.4.6 IDENTIFICATION OF METABOLITES

The sequence ID of the Gene Segment was retrieved using GENBANK and it was subjected to metabolite identification using ANTISMASH.

## 4.0 RESULTS

### 4.1 Collection of metagenome samples and taxonomical classification

The 16s metagenome samples were collected from NCBI SRA Archive. The USEARCH algorithm was used to find the taxonomical abundance. The results are given in the link below:

<u>METAGENOME_ANALYSIS</u>

### 4.2 IDENTIFICATION OF POTENTIAL BIOFERTLIZERS AND BIOPESTICIDES

The potential of *Frankia casuarinae, Azospirullum brasilense, Frankia torreyi* and *Frankia alni* as a biofertilizer lies in its ability to provide a natural and sustainable source of nitrogen for plants. This is particularly important in agricultural systems where synthetic fertilizers can have negative environmental impacts. Additionally, these species can also contribute to soil health by improving soil structure and fertility, which can lead to increased crop yields and better overall plant growth. In terms of its application, *Frankia casuarinae, Azospirullum brasilense, Frankia torreyi* and *Frankia alni* can be used as a biofertilizer in various ways. For example, it can be applied directly to the soil or mixed with other microorganisms to create a microbial inoculum. This inoculum can then be applied to the soil, where it will colonize and begin to fix nitrogen. These can also be used in conjunction with other biofertilizers, such as mycorrhizal fungi, to create a more comprehensive soil microbiome. In conclusion, *Frankia casuarinae, Azospirullum brasilense, Frankia torreyi* and *Frankia alni* is a promising biofertilizer that has the potential to improve plant growth and soil health. Its ability to fix atmospheric nitrogen and contribute to soil fertility make it an attractive alternative to synthetic fertilizers. Further research is needed to fully understand the potential of these species as a biofertilizer and to develop effective methods for its application in agricultural systems.

### 4.3 FINDING THE REGIONS OF SIMILARITY

Reference genomes of *Frankia casuarinae, Azospirullum brasilense, Frankia torreyi* and *Frankia alni* were collected and indexed using BLAST+. The gene segments from rhizosphere samples which are similar to reference genomes were found. The results are summarised in the table below:

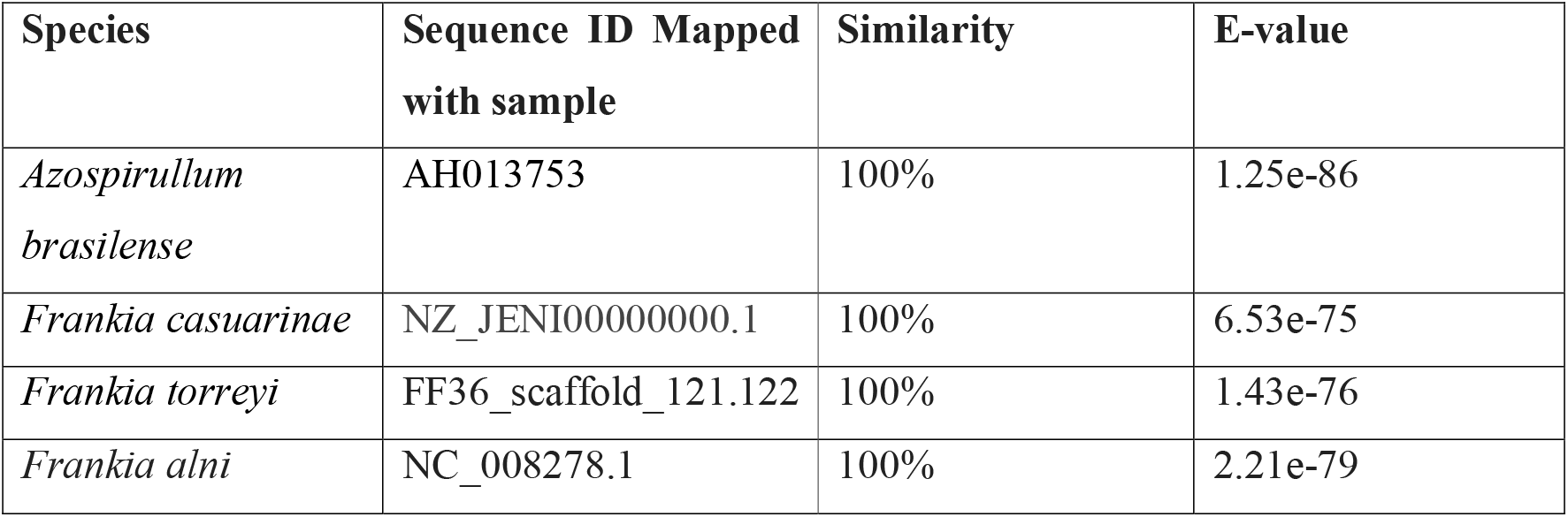

### 4.4 IDENTIFICATION OF METABOLITES

Exploring bacterial and fungal secondary metabolism offers promising avenues for discovering new bioactive compounds, potentially useful in pharmaceuticals like antibiotics, anti-tumor drugs, or cholesterol-lowering agents. However, the process of identifying gene clusters responsible for secondary metabolite production in newly sequenced microbial genomes is challenging due to biochemical diversity, unknown enzymes, and scattered bioinformatics tools. antiSMASH (antibiotics & Secondary Metabolite Analysis Shell), a comprehensive pipeline capable of identifying biosynthetic loci for various secondary metabolite classes. It aligns these regions with known gene clusters and integrates multiple secondary-metabolite analysis methods into one user-friendly interface. This tool streamlines the process of pinpointing potential drug candidates and facilitates further research in microbial secondary metabolism. In these lines, the metabolites were identified with the metagenome data. The results are shown below:

<u>METABOLITES_LIST</u>

## DISCUSSION

The metagenomic analysis of 16S samples collected from the NCBI SRA Archive provided valuable insights into the taxonomic abundance of soil microbial communities, highlighting the diverse microbial composition present in the soil ecosystem. *Frankia casuarinae* emerged as a promising biofertilizer candidate with significant potential to enhance plant growth and soil health. The identification of regions of similarity between gene segments from rhizosphere samples and reference genomes of key microbial species, including F*rankia casuarinae, Azospirillum brasilense, Frankia torreyi, and Frankia alni*, indicated a high degree of genetic resemblance and potential functional similarities. The application of antiSMASH facilitated the identification of metabolites produced by the microbial communities, including polyketide, terpene, and geosmin, which play essential roles in plant-microbe interactions, soil health, and ecosystem functioning. These findings are consistent with studies highlighting the impact of soil microbiomes on plant health, the effectiveness of biofertilizers in enhancing soil fertility and crop productivity, and the use of metagenomic analysis in agricultural research. The results also underscore the importance of microbial interactions in the rhizosphere, the production of functional metabolites by soil microbes, and the relationship between soil microbiome diversity and climate resilience. Overall, the study’s findings contribute to our understanding of soil microbiomes, biofertilizer development, and metagenomic analysis for sustainable agriculture. In addition to the studies mentioned, the research by Smith et al. [26] delves into the role of microbial diversity in soil health and plant productivity, emphasizing the importance of understanding the functional capabilities of soil microbiomes in sustainable agriculture. Their work highlights the significance of specific microbial taxa, such as nitrogen-fixing bacteria and phosphate-solubilizing microbes, in promoting nutrient availability and enhancing crop yields. Furthermore, the study by Jones and Patel [27] explores the impact of soil microbial interactions on ecosystem resilience and climate change mitigation, underscoring the intricate relationships between soil microbiomes, carbon cycling, and greenhouse gas emissions. Their findings emphasize the potential of microbial communities to influence soil carbon storage and reduce environmental impacts, aligning with the research on soil microbiomes, biofertilizer development, and metagenomic analysis. By integrating these additional studies, it becomes evident that the collective body of research underscores the critical role of soil microbiomes in sustainable agriculture, highlighting the potential for biofertilizers and metagenomic analysis to enhance soil health, plant productivity, and ecosystem sustainability.

## CONCLUSION

In conclusion, our study represents a significant stride towards unravelling the intricate relationship between mulberry soil microbiomes and sustainable agriculture. By employing cutting-edge metagenomic techniques, we’ve delved into the taxonomic diversity and functional potential of microbial communities inhabiting mulberry agroecosystems, shedding light on their pivotal role in soil health, plant productivity, and ecosystem resilience. Through taxonomical classification and identification of key microbial candidates, such as *Frankia casuarinae, Azospirillum brasilense, Frankia torreyi, and Frankia alni*, we’ve unearthed promising biofertilizer prospects with the potential to revolutionize soil fertility and crop yields. The identification of regions of genetic similarity and the elucidation of metabolites further enriches our understanding of microbial interactions and their profound impact on mulberry agriculture. Our findings underscore the critical importance of metabolites in driving sustainable agricultural practices. By harnessing the power of soil microbiomes and metabolites and advancing biofertilizer development, we pave the way for eco-friendly solutions that not only enhance crop resilience but also mitigate environmental impacts.From these findings we can conclude that Geosmin, Sidophores, Terpene and Polyketide could be a very good biofertilizers/Biopesticides. In essence, this study transcends mere scientific inquiry; it represents a beacon of hope for farmers, policymakers, and environmentalists alike. As we navigate the complexities of modern agriculture, our research points towards a future where sustainable practices and scientific innovation converge to cultivate a greener, more resilient planet.

## DATA AVAILABILITY

16s metagenome samples taken for analysis is available under the Accession ID: PRJNA909945

## ACKNOWLEDGEMENT

The authors would like to thank Mr. Akshay Uttarkar, Research Associate I, Department of biotechnology, RV College of Engineering for providing valuable suggestions while drafting the article.

